# LTRs activated by Epstein-Barr virus-induced transformation of B cells alter the transcriptome

**DOI:** 10.1101/233163

**Authors:** Amy Leung, Candi Trac, Hiroyuki Kato, Kevin R Costello, Rama Natarajan, Dustin E Schones

**Author notes:** **Corresponding Authors**: Amy Leung, Dustin E. Schones. Amy Leung, Ph.D., Department of Diabetes Complications and Metabolism, City of Hope, Duarte, CA, 91016. Dustin Schones, Ph.D., Department of Diabetes Complications and Metabolism, City of Hope, Duarte, CA, 91016.

## Abstract

Endogenous retroviruses (ERVs) are ancient viral elements that have accumulated in the genome through retrotransposition events. Although they have lost their ability to transpose, many of the long terminal repeats (LTRs) that originally flanked full-length ERVs maintain the ability to regulate transcription. While these elements are typically repressed in somatic cells, when this repression is lost, they can function as transcriptional enhancers and promoters. The mechanisms driving LTR activation, however, are not well understood. Epstein-Barr virus (EBV), which transforms primary B cells into continuously proliferating cells, is a tumor virus associated with lymphomas. We report here that transformation of primary B cells by EBV leads to genome-wide activation of LTR enhancers and promoters. The activation of LTRs coincides with local DNA hypomethylation and binding by transcription factors such as RUNX3, EBF1, and EBNA2. The set of EBV-activated LTRs is unique to transformed B cells when compared to other cell lines known to have activated LTRs. Furthermore, we found that EBV-induced LTR activation impacts the B cell transcriptome by upregulating transcripts driven by cryptic LTR promoters. These transcripts include genes important to oncogenesis of Hodgkin lymphoma, as well as those found in other cancers such as *HUWE1/HECTH9*. These data suggest that the activation of LTRs by EBV may be important to the pathology of EBV-associated cancers. Altogether, our results indicate that EBV-induced transformation of B cells alters endogenous retroviral element activity, thereby impacting host gene regulatory networks and oncogenic potential.

## Introduction

Endogenous retroviruses (ERVs) make up ˜8% of the human genome (Lander et al. 2001). A complete ERV locus contains two long terminal repeats (LTRs) flanking the Gag-, Poland Env-coding sequences needed for transposition activity (Mager and Stoye 2015). Active ERVs can transpose to new locations in the genome through RNA intermediates transcribed by RNA polymerase II (Stoye 2012). The transcription of ERVs is driven by LTRs which contain both enhancer and promoter elements (Stoye 2012). Most LTR loci are repressed by DNA methylation and histone modifications to inhibit their activity in the host genome (Wolf and Goff 2008; Rowe and Trono 2011; Friedli et al. 2014; Liu et al. 2014). In the human genome, the vast majority of ERVs are solitary LTRs, and thus incapable of transposition, but many still maintain the ability to enhance and promote transcription under certain conditions. A subset of these LTRs have been co-opted in the genome as cryptic enhancers and promoters for transcription (Maksakova et al. 2008; Rowe and Trono 2011; Leung and Lorincz 2012; Thompson et al. 2016). This LTR-driven transcription occurs in a cell- and tissue-specific manner (Faulkner et al. 2009).

There are many examples in which LTR-driven transcripts, both protein-coding and non-coding, play a role in pluripotent cells. In the early embryo of both humans and mice, specific ERV groups are transcriptionally active and drive expression of many transcripts in the developing cells (Evsikov et al. 2004; Peaston et al. 2004; Macfarlan et al. 2012; Maksakova et al. 2013; Göke et al. 2015). Likewise, both embryonic stem cells and induced pluripotent stem cells express LTR-driven transcripts, suggesting that they are important for pluripotency (Leung and Lorincz 2012; Fort et al. 2014; Lock et al. 2014; Lu et al. 2014; Ohnuki et al. 2014; Hashimoto et al. 2015; Babaian et al. 2016; Klawitter et al. 2016). In pathways other than development, LTR-driven transcripts have also been found to be expressed in autoimmune diseases and several types of cancer, including lymphoma, hepatocellular carcinoma, and prostate cancer (Lamprecht et al. 2010; Prensner et al. 2013; Lock et al. 2014; Hashimoto et al. 2015; Babaian et al. 2016; Chuong et al. 2016). While this work has led to an appreciation of the degree of aberrant LTR activation in normal and disease cells, the mechanisms that lead to LTR activation have remained unclear.

Originally discovered in cells derived from a Burkitt’s lymphoma biopsy, Epstein-Barr virus (EBV) can readily infect primary B lymphocytes (B cells) in vitro and in vivo and transform them into continuously proliferating lymphoblastoid cell lines (LCLs) in vitro, leading to the classification of EBV as a tumor virus (Epstein et al. 1964; Pope et al. 1973). In addition to lymphomas, EBV has been associated with many other cancer types, including nasopharyngeal carcinoma, gastric adenocarcinoma, and lymphoepithelioma-like carcinomas (Hsu and Glaser 2000). EBV can infect B cells and epithelial cells (Sixbey et al. 1984; Walling et al. 2001; Pegtel et al. 2004). In memory B cells, EBV can remain quiescent in a latent state using one of four latency programs, all of which are characterized by distinct expression of EBV-encoded genes, including transcription factors (Thorley-Lawson 2001). Transformation of primary B cells leads to genome-wide transcriptional changes including a decrease of apoptotic genes and an increase of proliferative genes (Allday 2013; Price and Luftig 2014; Price et al. 2017). EBV-induced transformation of primary B cells also leads to DNA hypomethylation at about two-thirds of the genome, similar to what is observed in cancer cells (Hernando et al. 2013; Hansen et al. 2014). It has furthermore been found that EBV infection in primary B cells transactivates a human ERV locus, HERV-K18, leading to the expression of a superantigen important for T cell response (Sutkowski et al. 1996).

To evaluate the impact of EBV infection of primary B cells on LTRs across the genome, we profiled H3K4me3, a chromatin modification associated with both active enhancers and promoters, in donor matched primary B cells and LCLs. We report here that LCLs have widespread activation of LTRs compared to primary B cells. Cryptic LTR activation as enhancers and promoters is coincident with loss of DNA methylation in LCLs at LTR loci and binding by B cell transcription factors (TFs), and EBNA2, a transcription factor encoded by the viral genome. A consequence of cryptic LTR activation is the expression of a number of chimeric (LTR-driven) mRNAs and non-coding RNAs originating from LTR promoters. We found that EBV transformation is sufficient to promote the expression of LTR-driven chimeric transcripts for *CSF1R* and *IRF5*, which are expressed in lymphomas but not in primary B cells and are important for growth and survival of Hodgkin Lymphoma cells. Additionally, we discovered a previously unannotated cryptic LTR-driven chimeric transcript for *HUWE1/HECTH9*, which encodes an E3 ligase that regulates the transcriptional activity of MYC and known to be important in blood cancers. Altogether, this suggests that LTR activation induced by EBV may support expression of key genes related to cancer development.

## Results

### H3K4me3 profiling reveals widespread enrichment of the modification at LTRs in LCLs

To explore the potential impact of EBV-transformation on the chromatin environment of B cells, we generated H3K4me3 profiles from three donor primary B cells and matched LCLs derived from the same donor B cells (**Fig. 1A; Supplemental Fig. S1A**). We identified a union set of 105,967 peaks (sites) of H3K4me3, a chromatin modification associated with enhancers and promoters, from B cells and LCLs. To identify genes that are differentially enriched for H3K4me3, we performed DESeq2 analysis (**Methods and Supplemental Methods**) on read counts found in the 2kb region surrounding the promoter (i.e. +/- 1kb) of genes (hg19 RefSeq annotation) that overlap a H3K4me3 site and found 9,654 sites as variable across the two groups (adjusted *P <0.01*) with 5,853 increased and 3,801 decreased (**Fig. 1B**, *top*). Upregulated sites were enriched at promoters of genes with the top KEGG term of ‘Pathways in cancer’ (**Supplemental Fig. S1B**). Genes found in this KEGG term included many genes known to be upregulated by EBV infection, including *FAS, IKBKB, and TRAF1* (**Supplemental Fig. S1C**) (Hernando et al. 2013). These data demonstrate that the H3K4me3 profiles of primary B cells and LCLs capture known transcriptional changes at genes related to the transformation of B cells to LCLs.

**Figure 1:**
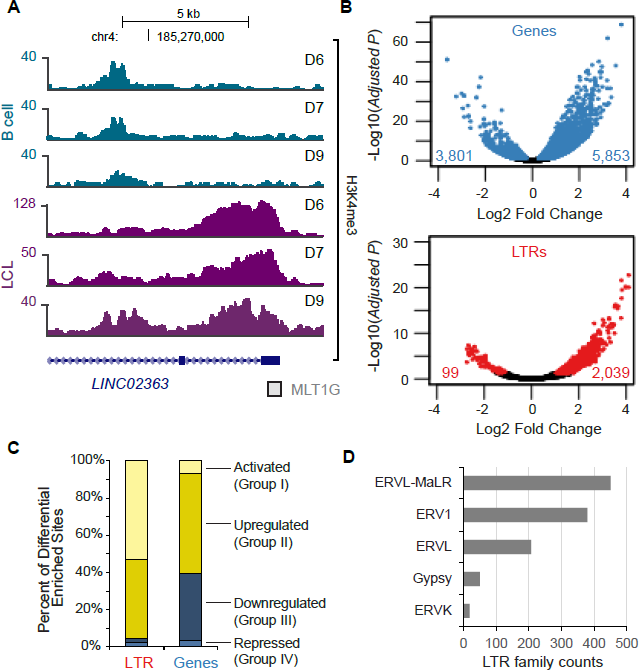
**EBV-induced transformation of B cells activates LTRs.** (*A*) Genome browser tracs of H3K4me3 profiles at *LINC02363*, a long non-coding gene, with the MLT1G LTR at the transcriptional start site. (*B*). Volcano plots of normalized H3K4me3 read counts in LCLs relative to B cells at (top) transcriptional start sites of genes and (bottom) at LTRs. (*C*) Percentage of differentially enriched H3K4me3 LTRs and genes classified as “Activated”, “Upregulated”, “Downregulated”, and “Repressed”. (*D*) Enumeration of the LTR families that are activated in LCLs (Group I).

We next examined the possibility that EBV-induced transformation influences transcriptional activity, as measured by increase in H3K4me3 enrichment, of LTR promoters and enhancers across the genome. Taking all LTR sites (RepeatMasker (Smit et al. 2013-2015)) found in the human genome, we performed DESeq2 analysis with read counts mapping to LTR sites that overlap a H3K4me3 site. Of the 2,138 differentially enriched LTR sites (adjusted *P <0.05*) discovered by this analysis, the vast majority (2,039) display increase in H3K4me3 enrichment in LCLs compared to B cells (**Fig. 1B**, *bottom*). These LTRs sites can be found at the transcriptional start sites (TSSs) of both long non-coding RNAs (e.g. *LINC02363*, **Fig. 1A**) and protein-coding genes (**Supplemental Fig. S2)**. In comparison to TSSs of genes, which show roughly equivalent degrees of increase and decrease in H3K4me3 enrichment in LCLs compared to B cells (**Fig. 1B**, *top*), LTRs have a marked increase in H3K4me3 in the LCLs (**Fig. 1B**, *bottom*). We next distinguished the LTRs and genes that are: Group I: “Activated” by complete gain of H3K4me3 (i.e. not enriched for H3K4me3 in B cells), Group II: “Upregulated” by increase in H3K4me3 (i.e. enriched with H3K4me3 in B cells), Group III: “Downregulated” (i.e. decrease in H3K4me3, but still enriched in LCLs, and Group IV: “Repressed” (i.e. no H3K4me3 enrichment in LCLs). With this classification, 1,134 LTRs are found in to be in Group I, and from hereon will be referred to as “activated LTRs”. These data indicate a widespread activation of LTR promoters after EBV transformation. HERV-K18, an ERV1 element known to be induced by EBV infection is found in the activated group, as expected (**Supplemental Fig. S3**). Examining the specific subfamilies of LTR promoters that are activated by EBV transformation revealed that the majority of the activated LTRs belong to either the ERV1 or ERVL-MaLR/ERVL families of LTRs (**Fig. 1D**). Interestingly, while there is great variety in the subfamilies of LTRs (241 subfamiles) (**Supplemental Table S1**), we found a dramatic activation of MLT subfamily members than expected by chance (e.g. MLT1D (*P* = 0.013), MLT1K (*P* =0.008), and MLT1F1 (*P* = 0.002)), and MER41 subfamily members (e.g. MER41A (*P* < 0.001, and MER41B (*P =* 0.011) (**Supplemental Table S1**). This data indicates that specific subfamilies of LTRs may be more susceptible to activation. To assess if there are any DNA rearrangements at LTRs that may lead to activation of these LTRs, we performed PCR analysis of genomic DNA and confirmed that there is no obvious rearrangements in the two assayed LTRs (**Supplemental Fig. S4**).

### Cryptic LTR promoter and enhancer activation coincides with loss of DNA methylation and binding of transcription factors

LTRs can be silenced through repressive histone modifications (Rowe and Trono 2011; Leung and Lorincz 2012; Liu et al. 2014), co-repressors (Jacobs et al. 2014; Wolf et al. 2015) and DNA methylation (Wolf and Goff 2008). EBV transformation of B cells has been shown to induce large-scale loss of DNA methylation, akin to the loss of DNA methylation in cancer cells (Hernando et al. 2013; Hansen et al. 2014). We therefore examined the DNA methylation profiles of activated LTRs after EBV transformation. Utilizing whole genome bisulfite sequencing data from primary B cells and matched LCLs (GSE49627) (Hansen et al. 2014), we examined DNA methylation and H3K4me3 (this paper) at the activated LTRs. Comparing the average CpG methylation level change at LTRs in three donor B cells with the three LCLs revealed that activated LTRs are generally hypomethylated upon EBV treatment and have corresponding increases in H3K4me3 (**Fig. 2A,** representative example **Fig. 2B**). In contrast to the large-scale hypomethylated blocks previously reported in EBV induced B cell immortalization, the DNA methylation changes we observed were more local changes, specific to the LTR elements (**Fig. 2B, Supplementary Fig. S5**). This data supports the idea that EBV-induced hypomethylation is concurrent with LTR activation. We next addressed whether loss of DNA methylation itself is sufficient to activate LTRs. Examining DNA methylation at all LTRs, regardless of EBV-activation status, indicated that hypomethylation does not necessarily result in gain of H3K4me3 modifications after EBV transformation (**Fig. 2C**), suggesting that loss of DNA methylation is not sufficient to activate LTRs in B cells and that additional regulatory factors are required.

**Figure 2:**
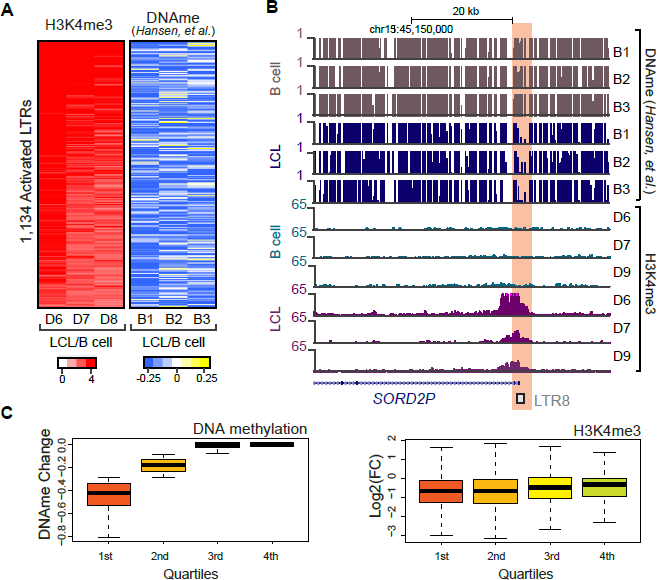
**Activated LTRs are hypomethylated in LCLs.** (*A*) (left) Heatmap of H3K4me3 in three B cell donors and three matched LCLs at activated LTRs and (right) heatmap of DNA methylation changes in matched B cells and LCLs at activated LTRs. (*B*) Genome browser tracks of DNA methylation and H3K4me3 profile in B cells and LCLs at *SORD2P* locus driven by an LTR8. (*C*) (left) Boxplot of DNA methylation changes and (right) H3K4me3 change between B cells and LCLs stratified into quartiles based on DNA methylation change.

To gain a better understanding of the regulatory factors involved in activating LTRs, we next examined transcription factor binding at activated LTRs in GM12878 cells using ChIP-seq data for TFs generated from ENCODE (Dunham et al. 2012), along with EBNA2 (GSE29498), RPBJ (GSE29498), and NF-κB subunits (GSE55105) (Zhao et al. 2011; Zhao et al. 2014). Overall, we found that there are 36 transcription factors bound to at least 25% of the activated LTR sites (**Fig. 3A**). A number of these transcription factors, as expected based upon existing literature, are upregulated including RUNX3, EBF1, and EBNA2 (**Supplemental Fig. S6**). Strikingly, ˜67% of the LTRs activated by EBV are bound by RUNX3, a transcription factor expressed throughout the hematopoietic system that is specifically upregulated by EBV during LCL development and promotes cell proliferation (Spender et al. 2002; Brady et al. 2009). EBF1, which is important for B cell lineage development (Boller and Grosschedl 2014) is also among the most enriched transcription factors at activated LTRs. IKAROS/IKZF1, which binds to ˜25% of activated LTRs, plays key roles in the latent-lytic switch of EBV in B cells, indicating that perhaps LTR activation is influenced by EBV-induced pathways (Iempridee et al. 2014). Interestingly, EBNA2, a TF encoded in the EBV genome, is bound to ˜30% of the EBV-activated LTR (**Fig. 3A**), Given that many transcription factors bind to common regulatory regions (Wang et al. 2013), we examined the co-occupancy of transcription factors bound at activated LTRs by performing hierarchical clustering of pairwise correlations of binding patterns (Methods). This analysis revealed a number of clusters that appear to co-occupy LTRs together (**Fig. 3B**). EBNA2 co-binds with RBPJ, as has been previously demonstrated ((Zhao et al. 2011)). Other clusters include a RUNX3 cluster with many of the factors binding the largest fraction of activated LTRs, a general TF/chromatin cluster with p300/CHD2, and an NF-κB cluster with RELA, RELB, cREL and p52 (**Fig. 3B**). We next asked whether a particular subfamily is more likely to be bound by the top 10 transcription factors. Overall, the LTR subfamily bound by these transcription factors are diverse (**Supplemental Fig. S7**, **Supplemental Table S2**). We next examined the number of EBV-activated LTRs that give rise to transcripts using CAGE (Cap-analysis of gene expression) data from GM12878 (ENCODE) (Hoffman et al. 2013). In total, ˜31% of EBV-activated LTRs overlap CAGE enriched sites, indicating this subset of LTRs can initiate transcription (**Fig. 3C**). Those without CAGE evidence are likely either transcribing weakly or serving as regulatory regions for other genes.

**Figure 3:**
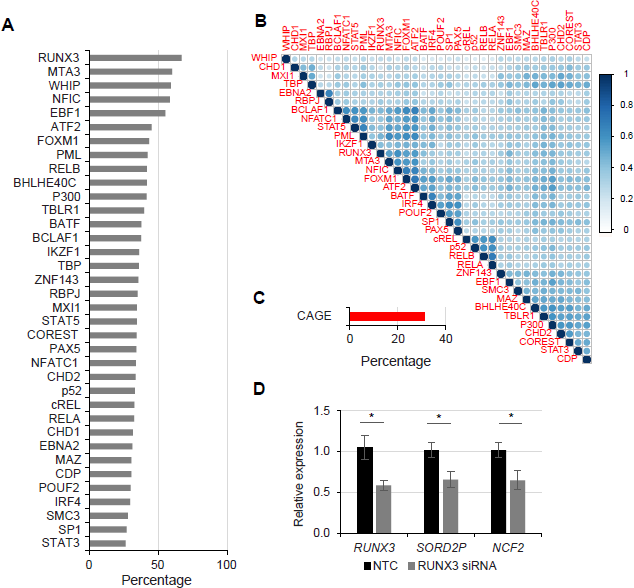
**Activated LTRs are binding sites for many transcription factors in GM12878 and can promote transcription.** (*A*) Bargraphs of the percentage of activated LTRs bound by each transcription factor. (*B*) Correlation plots (Pearson) of transcription factor co-occupancy at EBV-activated LTRs. Hierarchical clustering was performed to identify transcription factors that co-occur. (*C*) Percentage of activated LTRs with CAGE sequences in GM12878 indicating promoter activity. (*D*) RT-qPCR of *RUNX3*, *SORD2P*, and *NCF2* transcripts after siRNA knockdown of *RUNX3* transcripts (* denotes *P*<0.05).

To determine if RUNX3 binding at LTRs was sufficient to drive transcription of chimeric transcripts, we performed siRNA knockdown of RUNX3 and examined the expression of several genes driven by cryptic LTR promoters (**Fig. 3D**). Upon RUNX3 knockdown, the genes we examined were reduced in expression, consistent with RUNX3 binding at cryptic LTR promoters driving the expression of these genes.

### LTRs activated by EBV are mostly unique to LCLs

Given that LTRs have previously been shown to be active in other cells lines, including H1 (embryonic stem cell) and K562 (immortalized myelogenous leukemia) cells (Djebali et al. 2012; Fort et al. 2014), we next addressed whether the LTRs activated by EBV are unique to LCLs or are generally active in cell lines. Using H3K4me3 profiles generated by ENCODE from GM12878, H1, Hela S3, HUVEC, HepG2, and K562 cells (Dunham et al. 2012), we assessed the enrichment of H3K4me3 across these cells lines at the LTRs activated by EBV. Using hierarchical clustering of the H3K4me3 profiles generated from two different sequencing centers within the ENCODE project, we found that the profiles from the same cell lines cluster together and that the vast majority of the EBV-activated LTRs are not enriched for H3K4me3 in other cells (**Fig. 4A**). These data suggest that the activation of LTRs by EBV occurs at a subset of loci that are unique from those activated in other cells. Indeed, analysis of H3K4me3 enrichment at all LTRs across the six cells revealed that each cell line has a distinct profile of active LTRs (**Supplemental Fig. S8**). We further profiled H3K4me3 profiles in different normal tissue types from RoadMap data, and find that the enrichment at the activated LTRs is low compared to LCLs (**Supplemental Fig. S9**). This data agrees with previous reports that LTR expression and activity is cell type-specific (Faulkner et al. 2009; Jacques et al. 2013; Xie et al. 2013).

**Figure 4:**
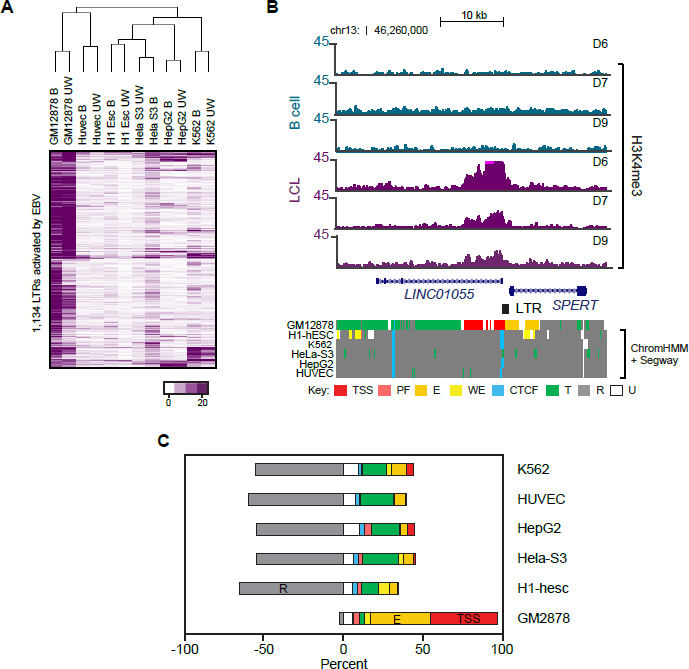
**Transcriptional activity of activated LTRs is mostly specific to transformed B cells.** (*A*) Hierarchical clustering of H3K4me3 read counts across cell liens from ENCODE data. (B: Broad; UW, University of Washington; replicates from different sequencing centers). (*B*) Genome browser track of h3K4me3 profile in B cells and LCLs, and of ChromHmm + Segway combined data at the *LINC01055-SPERT1* locus (Key: TSS, predicted promoter region including TSS; PF, predicted promoter flanking region; WE, predicted weak enhancer or open chromatin cis regulatory element; CTCF, CTCF enriched element; T, predicted transcribed region; R, predicted repressed or low activity region; U, unclassified). *(C*) ChromHmm + Segway combined classification of all activated LTRs shown as a frequency of total for each ENCODE cell line (color schemes as in (*B*)).

**Figure 5:**
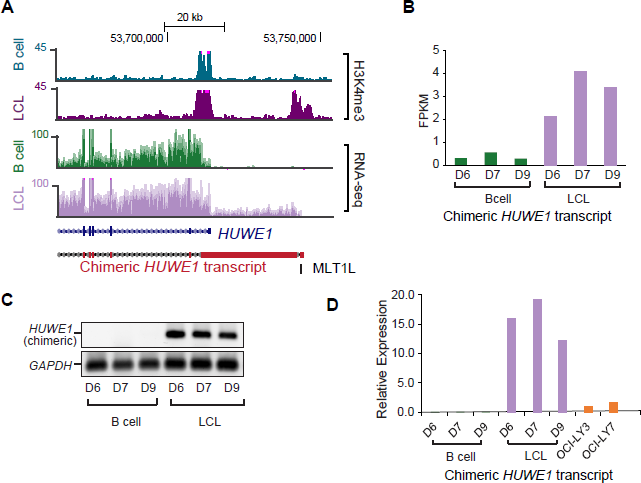
**Activated LTRs can drive transcription of chimeric transcripts relevant to cancer.** (*A*) Genome browser tracks for H3K4me3 and RNA-seq profile for B cells and LCLs at *HUWE1/HECTH9* locus. Shown in red is the chimeric transcript assembled from RNA-seq dataset. Transcriptional start site of alternative chimerical transcript overlaps MLT1L. (*C*) FPKM (fragments per kilobase per million reads) in each RNA dataset. (*D*) RT-PCR analysis of chimeric *HUWE1* transcripts in B cells and matched LCLs. (*E*) RT-qPCR analysis of chimeric *HUWE1* expression in B cells, LCLs, and two EBV-negative lymphoma cell lines.

We furthermore examined whether these EBV-activated LTRs in GM12878 are classified as transcriptionally active regulatory regions. At specific loci, such as at *LINC01055* (**Fig. 4B**), the LTR marked by H3K4me3 is classified by combined analysis from Segway and ChromHMM (Ernst and Kellis 2010; Ernst and Kellis 2012; Hoffman et al. 2012; Hoffman et al. 2013) as a TSS. Examining the functional classification from Segway and ChromHMM of activated LTRs across the genome, we found that the majority of the EBV-activated LTRs are classified as promoters or enhancers (>80). In contrast, these same regions are mostly classified as repressed regions in other cell lines (**Fig. 4C**).

### EBV-induced activation of LTRs is sufficient to drive lymphoma-related and cancer-related transcripts

Activation of cryptic LTR promoters has been demonstrated to promote expression of TE-driven transcripts (Babaian and Mager 2016; Elbarbary et al. 2016). To assess the potential impact of LTR activation, we examined whether EBV-activated LTRs can lead to expression of chimeric transcripts. By combining *de novo* assembly of transcripts and enrichment of H3K4me3 at the 5’ end (Methods), we found that there are 24 LTR-driven transcripts that are upregulated in the LCLs (**Supplemental Table S3**). These include the transcripts highlighted above, including *SORD2P* and *LINC02363*. Furthermore, expression of these TE-driven transcripts is highest in EBV-transformed B cells compared to all other tissues examined from GTEx data (**Supplemental Fig. S10**).

Of the LTR-driven chimeric transcripts identified in our dataset, several are relevant to lymphoma development. A THE1B LTR drives the expression of a non-canonical isoform of *CSF1R*, leading to lineage inappropriate expression of the gene in lymphoma cells (Lamprecht et al. 2010) and a LOR1a LTR drives the expression of *IRF5*, which is a key regulator in the development of lymphomas (Babaian et al. 2016). By RT-QPCR analysis of RNA isolated from B cells and LCLs, we found that the non-canonical transcript is indeed expressed in LCLs, but not in B cells (**Supplemental Fig. S11**). The canonical *CSF1R* transcript, however, is expressed in both LCLs and primary B cells, which has previously been observed (Baker et al. 1993). Similarly, in addition to the canonical transcript, LCLs express the non-canonical *IRF5* isoform driven by the LOR1a LTR (**Supplemental Fig. S11**). These data indicate that EBV-induced transformation of B cells is sufficient to activate LTRs driving transcripts known to be important in the development of lymphomas.

In addition to *CSF1R* and *IRF5*, the *HUWE1/HECTH9* locus has a chimeric/non-canonical isoform originating from an MLT1L LTR upstream from the canonical isoform (**Fig. 6A**). *HUWE1* is involved in the transcriptional activity of MYC and has been found to be essential to cell proliferation in cancer cells (Adhikary et al. 2005). From our RNA-seq dataset, expression of this chimeric *HUWE1* transcript is clearly upregulated in LCLs (**Fig. 6B**). To confirm the presence of this transcript, we performed RT-PCR specific to the isoform using primers unique to MLT1L and downstream exon 2 of *HUWE1* transcript. This analysis confirmed that the chimeric *HUWE1* transcript is expressed in the LCLs but not the B cells (**Fig. 6C**). We furthermore examined if the chimeric *HUWE1* transcript is specifically expressed in EBV transformed cells. Altogether, these data suggests that EBV-transformed LCLs specifically activates MLT1L at the *HUWE1/HECTH9* to express a chimeric isoform.

It has previously been shown that diffuse large B cell lymphoma (DLBCL) and Hodgekin lymphoma also has activation of TE promoters leading to chimeric transcript expression (Lock et al. 2014; Babaian et al. 2016). We examined our RNA-sequencing profiles to determine if the chimeric transcripts identified in DLBCL are also expressed in the LCLs. Comparing the datasets, we did not find common cryptic transcripts, other than *CSF1R* and *IRF5* suggesting that while LTR activation leading to chimeric transcripts occurs in both DLBCL and LCLs, the subset of chimeric transcripts is quite unique to EBV-induced transformed cells. To test this, we examined if the LTR-driven transcripts are unique to EBV-positive lines compared to EBV-negative lines. We analyzed the expression of five LTR-driven transcripts from the *HUWE1*, *LINC01055, CERNA2*, *SORD2P*, and *NCF2*, and found that the expression is higher in EBV-induced LCLs than in the EBV-negative lymphoma cell lines OCI-LY3 and OCI-LY7 (**Fig. 6D**, **Supplemental Fig. S12**). These data indicates that EBV-induced transformation of B cells drives specific activation of cryptic LTR promoters.

## Discussion

It is well established that transposable elements can alter gene regulatory networks in both normal and diseased states (Babaian and Mager 2016; Elbarbary et al. 2016). Previously, it has been shown that EBV infection of B cells drives expression of a HERV-K locus found within the intron of CD48 (Sutkowski et al. 1996; Sutkowski et al. 2001). This leads to expression of a superantigen proposed to be important for immune response to the mounting infection and perhaps clearance of the viral load (Sutkowski et al. 2001). Here, we report that activation of LTRs by EBV occurs pervasively across the genome from varying subfamilies of ERVs. The activation of LTRs involves both DNA hypomethylation and transcription factor binding. The transcriptional activity of EBV-activated LTRs is unique to transformed B cells compared to other cell types that are known to have active LTRs. Furthermore, we have shown that EBV-induced transformation of B cells leads to expression of many chimeric transcripts including several that are relevant to lymphoma biology, as well as other cancers types.

Viruses have previously been implicated in the widespread activation of LTRs and long interspersed nuclear elements (LINEs) in hepatocarcinomas, as tumors with viral etiology display a higher level of LTR and LINE activation (Shukla et al. 2013; Hashimoto et al. 2015; Schauer et al. 2018). Taken together with our data, this suggests that LTR activation is related to the host’s response to viruses. The most obvious pathway responsible for LTR activation are those related to immune response to viruses. Interestingly, ERVs have been evolutionarily co-opted by innate immune pathways (Chuong et al. 2016). Our current analysis of transcription factors bound to LTRs in GM12878 does find that NF-kB subunits can bind to ˜40% of LTRs suggesting these LTRs may have an important role in regulating the immune pathways. It has previously been demonstrated that the non-canonical isoform of *CSF1R* is regulated by an NF-κB binding site at a THE1B element (Lamprecht et al. 2010). Future analysis of the dynamics of transcription factor occupancy and signaling at LTRs across EBV infection time points will be necessary to fully elucidate this issue.

The most common transcription factor found at EBV-activated LTRs, bound to ˜67% of activated LTRs, is RUNX3. This transcription factor is highly upregulated by EBV-encoded transcription factors (Spender et al. 2002; Brady et al. 2009). Previously, RUNX3 has been reported to be involved in LINE-1 transcription (Yang et al. 2003). To our knowledge, this is the first report of RUNX3 involved in LTR activation. There has also been recent report that RUNX3 can induced site-specific DNA hypomethylation (Suzuki et al. 2017). Along with our finding that RUNX3 binds to >50% of all EBV-activated LTRs, EBV-induced RUNX3 activity may be an important factor in the hypomethylation of EBV-activated LTRs.

Global loss of DNA methylation is a hallmark of cancer cells. EBV-induced transformation is one exogenous stimulus that can lead to global loss of DNA methylation (Hansen et al. 2014). Our results indicate that LTR loci become hypomethylated upon EBV-induced transformation, and that a subset of these hypomethylated LTRs can then be transcriptionally activated. Interestingly, we found that DNA hypomethylation is insufficient to transcriptionally activate LTR loci. We furthermore found that many of these activated LTRs are bound by specific transcription factors, indicating that there may be both the need to derepress as well as activate by transcription factor binding. This derepression/activation model has been proposed previously by others (Lamprecht et al. 2010; Babaian and Mager 2016). This model suggests that while DNA hypomethylation can be achieved in different cells, the LTRs that are activated will be unique based upon the transcription factors that are present within the cell. In line with this model and previous data describing cell-type specific binding of transcription factors at transposons (Xie et al. 2013), our data show that LTR activation is a cell type-specific behavior.

With the previous observation that a HERV-K locus is expressed to promote superantigen expression (Sutkowski et al. 1996; Sutkowski et al. 2001), our finding that there is extensive LTR activation suggests that LTR activation by EBV may be an intrinsic process that the host cell has co-opted to benefit itself. Interestingly, the additional finding that these LTRs can lead to activation of *CSF1R* and *IRF5*, two genes important for lymphoma cell survival and proliferation, suggests that EBV activation of LTRs can also have a benefit for the virus in promoting survival of the infected cell. The activation of these two genes, however, may have,negative consequences for the host in the long run in the form of the oncogenesis. EBV has long been associated with increased risk of lymphoma (Young et al. 2016). Interestingly, analysis of lymphoma cell transcriptomes revealed many TE-driven chimeric transcripts (Lock et al. 2014; Babaian et al. 2016). Analysis of our transcriptome and chromatin profiles reveals additional LTR-driven transcripts including a chimeric isoform of *HUWE1/HECTH9*. This E3 ubiquitin ligase has previously been shown to be important in regulating the transcriptional activity of MYC, and as such may have an important role in LCL development as well as in the development of lymphomas related to EBV (Adhikary et al. 2005). Interestingly, compared to other lymphoma cell lines that are EBV-negative, this transcript and other LTR-driven transcripts in LCLs are highly expressed, suggesting that they may be a unique set that is expressed in EBV-positive cells. Additionally, transformed B cells only express a subset of those described in lymphoma, indicating that perhaps additional events that promote LTR activation occur in the development of lymphoma.

Based on our results, it is important to examine whether the subset of lymphomas, including Hodgkin lymphoma and diffuse large B-cell lymphoma (Lamprecht et al. 2010; Lock et al. 2014), that express LTR-driven transcripts, have been infected by EBV. We have preliminarily addressed this by examining the cell lines that express the non-canonical isoform of *IRF5* (Babaian et al. 2016) and found that it is expressed in both EBV-positive and EBV-negative B cell lymphoma cell lines tested by Babaian et al. This suggests that LTR activation does not solely rely upon viral activation, and that perhaps in the context of lymphoma development, activation of specific chimeric transcripts can occur through other means (other than viral activation).

The cell-type specificity of LTR activation may have important consequences. In transformed B cells, 24 chimeric transcripts are activated, and these have the potential to impact cellular behavior. Further analysis is needed to understand the potential function of these EBV-activated TE-driven transcripts. The location and arrangement of LTRs can govern the perhaps unique set of chimeric transcripts that can be expressed in any given cell type. These LTR-driven transcripts may be uniquely expressed in a particular disease state and therefore used as biomarkers of disease or unique therapeutic targets.

## Methods

### Collection of B cells and growth of lymphoblastoid cell lines

Blood samples from de-identified healthy donors were obtained following institutional guidelines at the City of Hope (IRB# 17022). B cell population was isolated using Dynabeads CD19 Pan B magnetic beads (Invitrogen). Isolated B cells were collected for chromatin immunoprecipitation, DNA isolation, and RNA isolation. LCLs were derived from B cells isolated from primary B cells (from de-identified donors labeled D6, D7, and D9) using EBV collected from B95-8 strain containing marmoset cell line (Hui-Yuen et al. 2011). Briefly, B cells were incubated with EBV-containing supernatant, grown for >6 weeks, frozen, and regrown to establish continual proliferating cell lines.

### Chromatin immunoprecipitation (ChIP) sequencing

In B cells and LCLs from donors, ChIPs were carried out with anti-H3K4me3 antibody (Abcam, ab8580) using standard ChIP protocols. Sequencing libraries were generated using the Kapa Hyper Prep kit (Kapa Biosystems) and sequenced using Illumina HiSeq 2500 technology. We obtained on average ˜65M 100bp x 100bp paired-end reads per H3K4me3 library ˜21M 100bp x 100bp paired-end reads per input library (H3K4me3, **Supplemental Table S4**). Sequences were aligned to the hg19 reference genome using Bowtie (Langmead 2010) with default options, except only uniquely mapped reads were retained (-m 1 option). BEDtools (Quinlan and Hall 2010; Quinlan 2014) was used to generate wiggle tracks which were visualized in the UCSC Genome Browser. Peaks of H3K4me3 were analyzed using MACS (Zhang et al. 2008). To identify H3K4me3 differentially enriched at genes, counts at peaks overlapping RefSeq transcriptional start sites were analyzed using comparing B cells to LCLs (Love et al. 2014). To identify H3K4me3 differentially enriched LTRs, all LTR loci (RepeatMasker Open-4.0) (Smit et al. 2013-2015) overlapping peaks were obtained and counts of H3K4me3 reads were analyzed using DESeq2. For example of commands, see **Supplemental Methods**.

For gene ontology analysis, differentially enriched H3K4me3 peaks at transcriptional start sites (RefSeq annotation, hg19) were analyzed. Genes with the transcriptional start sites were used as the list using DAVID Functional annotation Tool (Huang et al. 2009a; Huang et al. 2009b).

For EBNA2 and RPBJ binding sites, we downloaded ChIP-sequencing data for EBNA2 and RBPJ from GEO (GSE29498)(Zhao et al. 2011) and aligned the fastq data using Bowtie as with H3K4me3 data above. Peaks of enrichment were determined using MACS with default parameters.

### RNA-sequencing

RNA was extracted from B cells and LCLs using Trizol (Invitrogen). RNAs were depleted of ribosomal RNA (Ribo-Zero rRNA removal kit, Illumina). Eluted RNAs were prepared for sequencing according to Illumina protocols, and sequencedon the HiSeq 2500 Illumina platform. Reads were aligned to hg19 reference genome with hg19 Refseq gene annotation as a guide using HISAT with default parameteres (Kim et al. 2015). 56-65M 100bp x 100bp paired end reads were aligned for each library. Transcript assembles were built from aligned BAM files, filtered for redundant reads, using StringTie (Pertea et al. 2015) and guided using the Refseq transcript assembles for hg19 with default settings. Files were then merged using StringTie – merge guided using the Refseq transcript assembles for hg19 using default settings. Overlapping predicted transcripts were then merged using a custom python script and the 5’ ends of the transcripts were then intersected with the hg19 RepeatMasker annotation (RepeatMasker (Smit et al. 2013-2015)). LTR-driven transcripts mapping known internal exons were removed from the analysis. As an additional measure of bonafide transcripts originating from LTRs, only LTR-driven transcripts with 5’ ends that mapped with H3K4me3 sites were kept. Reads mapping to LTR-driven transcripts were then called using BEDtools coverageBed (Quinlan and Hall 2010; Quinlan 2014). Differential expression was detected using the EdgeR package (Robinson et al. 2010).

### Gene expression with PCR and qPCR analysis

RNA was isolated from donor B cells and LCLs using Trizol (Life Technologies). cDNA was generated from isolated RNAs using SuperScript IV Reverse Transcriptase Kit (Invitrogen) with random primers. PCR (30 cycles) was performed for *GAPDH*, and isoforms of *CSF1R, IRF5*, and *HUWE1*. For qPCR analysis, qPCR primers were generated for transcripts from *SORD2P*, *LINC01055*, *CERNA2*, *NCF2*, *RUNX3*, and *HUWE1* and used with KAPA SYBR Master Mix. Primers are available in **Supplemental Table S5.**

### Whole-genome bisulfite sequencing (WGBS) and bisulfite Sanger sequencing

To assess the changes in DNA methylation genome-wide, we obtained WGBS data of B cells and donor-matched LCLs from GEO database (GSE49629) (Hansen et al. 2014). BS-Seeker2 (Guo et al. 2013) was used to obtain DNA methylation levels genome-wide. To obtain changes in DNA methylation, individual CpG methylation levels overlapping LTRs were obtained and averaged across the LTR locus and the values were compared B cells to matched LCLs. For analysis of +/- 20kb regions surrounding LTRs, 500bp non-overlapping bins were generated +/-20kb. Averaged CpG methylation levels within each bin were determined and relative change was determined between matched B cell and LCL. To examine the impact of DNA methylation on changes in H3K4me3 at LTRs, averaged change in CpG methylation was determined for each LTR between matched B cell and LCLs. LTRs were stratified into quartiles and H3K4me3 fold change was determined for each quartile set of LTRs.

### ENCODE transcription factor, chromatin, and gene expression analysis

For analysis of transcription factor binding at LTRs, ENCODE data for transcription factor ChIP-sequencing from GM12878 (Dunham et al. 2012) were downloaded from UCSC Genome Browser, EBNA2 (GSE29498), RBPJ (GSE29498), and NF-kB subunits (GSE55105) were obtained. We surveyed ˜100 transcription factors. To determine occupancy of transcription factors at EBV-activated LTRs, presence or absence of transcription factor binding site from ENCODE at EBV-activated LTR was determine using BEDtools. For co-occupancy analysis, hierarchical clustering of pairwise correlations (Pearson) was performed.

To compare H3K4me3 enrichment at EBV-activated LTRs in GM12878 and other cell lines (H1, Hela S3, HepG2, HUVEC, and K562), H3K4me3 ChIP-seq fastq data was downloaded from UCSC Genome Browser. Reads were aligned to hg19 genome as described above using Bowtie. Number of reads (counts) was determined for each cell line and hierarchical clustering was performed across each LTR and across the cell lines. For the genome segmentation analysis, Combined Segway and ChromHMM data (Ernst and Kellis 2010; Ernst and Kellis 2012; Hoffman et al. 2012; Hoffman et al. 2013) were downloaded from UCSC Genome Browser. For each EBV-activated LTR, overlapping chromatin state was determined. For those with two or more overlapping chromatin states, the state with the greatest portion of the LTR was selected for frequency analysis.

To determine the impact of EBV-activated LTRs on neighboring genes in GM12878 and other ENCODE cell lines, we downloaded ENCODE gene expression data from UCSC Genome Browser (http://genome.ucsc.edu/encode/downloads.html). The immediate adjacent gene from EBV-activated LTR was determined and FPKM of that gene was analyzed across all cell lines.

### PCR analysis

Genomic DNA was isolated from B cells and LCLs with 0.5% SDS lysis buffer (0.5% SDS, 50mM HEPES pH 7.5, 150mM NaCl, 1mM EDTA,1.0% Triton X-100, 5mM NaF) and DNA was extracted with a standard phenol-chloroform/ethanol precipitation protocol. PCR analysis was performed with specific primers targeting HERSVS71 and LTR8. Primer sequences can be found in **Supplemental Table S5**.

### Transfections and knockdown of RUNX3

siRNA knockdown of *RUNX3* transcripts was performed with 1 million lymphoblastoid cells (D7 LCL) according to manufacturer’s protocol (Amaxa) using 50nM pool of siRNA targeting RUNX3 (Thermo Scientific, cat #AM16708; ID: 107390 and 115509) or 50nM of control siRNA (Non-targeting control, Thermo Scientific). Cells were recovered after transfections and grown for two days before collection of RNA analysis. Total RNA was extracted, cDNA was prepared as described above, and used for qPCR analysis.

### Data Access

All new sequencing data generated from this study is deposited to NCBI GEO repository (GSE106851).

## Acknowledgments

This work was supported by K01DK104993 (AL); R01HL106089, R01HL087864 and RO1DK065073 (RN); R01DK112041 and R01CA220693 (DES). Research reported in this publication included work performed in the Pathology and Integrative Genomics Cores supported by the National Cancer Institute of the National Institutes of Health under award number P30CA33572. We would like to thank Drs. Sandrine Lacoste and Timothy O’Connor for their helpful assistance with EBV transformations, and Dr. John Chan for sharing critical reagents with us.

